# Parameter-Dependent Effects of Spinal Cord Stimulation on Neural Activation and Evoked Compound Action Potentials

**DOI:** 10.64898/2026.08.03.742575

**Authors:** Meagan K. Brucker-Hahn, Hans J. Zander, David A. Dinsmoor, Scott F. Lempka

**Affiliations:** Department of Biomedical Engineering, University of Michigan, Ann Arbor, MI, USA; Biointerfaces Institute, University of Michigan, Ann Arbor, MI, USA; Medtronic plc, Minneapolis, MN, United States of America; Department of Anesthesiology, University of Michigan, Ann Arbor, MI, USA

**Keywords:** Chronic pain, Computational modeling, Evoked potentials, Spinal cord, Spinal cord stimulation

## Abstract

**Objective:** Spinal evoked compound action potentials (ECAPs) provide a quantitative measure of the neural response during spinal cord stimulation (SCS) and can be leveraged in closed-loop applications to control dose in response to spinal cord movement. However, interpretation of ECAPs recorded *in vivo* is limited by susceptibility to noise, inter-subject variability, and other confounding factors. As SCS systems evolve and the clinical and research applications of ECAPs expand, it is critical to understand how physiological and technical factors influence ECAP generation and morphology.

**Approach:** We used a computational modeling framework to systematically investigate the influence of anatomical (e.g., dorsal cerebrospinal fluid (dCSF) thickness), stimulation (e.g., pulse width, waveform shape, stimulation configuration, stimulation frequency), and recording configurations on the neural responses and ECAPs generated during SCS. We employed a hybrid computational modeling approach, coupling finite element method models with multicompartment axon models to simulate neural responses to SCS. Using these models, we characterized the spatiotemporal dynamics of neural recruitment and the resulting ECAP waveforms.

**Main results:** Neural responses and model ECAPs were strongly influenced by factors, such as dCSF thickness, pulse width, and stimulation waveform shape. Stimulation parameters introduced trade-offs between axonal recruitment thresholds, neural activation selectivity, and ECAP timing and morphology. Notably, similar ECAP amplitudes could obscure differences in the underlying neural recruitment. Complex ECAP morphologies also emerged in response to distinct stimulation paradigms, reflecting changes in the spatiotemporal properties of axonal activation. Additionally, we demonstrate that the selection of recording electrodes can be optimized to enhance recorded ECAP amplitudes.

**Significance:** Our findings provide a theoretical framework to advance our mechanistic understanding of SCS-induced ECAPs and offer insights into optimization strategies to improve closed-loop SCS therapies.

## 1. Introduction

Spinal cord stimulation (SCS) is a widely used neurostimulation therapy for the treatment of chronic pain. Over the past 50 years [1,2], innovations in SCS have significantly expanded the range and complexity of available stimulation parameters. The clinical efficacy of SCS is highly dependent on appropriate parameter selection, which remains a complex and individualized process [3–7]. More recently, closed-loop SCS paradigms have emerged which leverage the ability to record stimulation-evoked neural responses, termed evoked compound action potentials (ECAPs), as real-time control signals to automatically adjust stimulation [8–10]. These systems have demonstrated improved patient outcomes compared to traditional open-loop approaches by maintaining more consistent neural activation to mitigate over- or under-stimulation [8–10].

Beyond their clinical utility, ECAPs have offered valuable insights into the mechanisms of SCS [11–22]. For example, ECAP characteristics vary with stimulation parameters and anatomical factors, providing insights into fundamental aspects of neural activation during SCS. These variations can be utilized to infer properties of neural recruitment that would otherwise be inaccessible in clinical and experimental settings. However, ECAP interpretation is complicated by multiple technical and physiological factors. Recorded ECAPs are highly susceptible to physiological and electrical noise [12,17,23–25]. Additionally, in clinical and experimental studies, stimulation parameters and anatomical variables are often unintentionally co-varied, making it difficult to isolate the effects of individual variables. For example, examining the effect of a stimulation waveform may be confounded by differences in lead placement or subject-specific anatomy, limiting the generalization of findings [13,14]. As ECAP-based closed-loop SCS systems continue to be used clinically and their application expands beyond conventional tonic stimulation [26–28], it is essential that we understand the factors that influence ECAP generation and morphology.

Computational models serve as a powerful tool for systematically investigating the neural response to SCS [11,15,29–34]. Unlike clinical and experimental studies, computational models provide complete control over stimulation and anatomical variables, eliminating confounding factors such as inter-subject variability and stimulation artifact. Therefore, in this study, we used a computational modeling framework to characterize how individual variations in stimulation parameters, recording configurations, and anatomy influence neural responses to SCS that are reflected in ECAP recordings. We combined a finite element method (FEM) model of the spinal cord, surrounding anatomy, and implanted electrode array with multicompartment axon models to simulate the neural response to SCS. We calculated model ECAPs using a reciprocal approach to estimate the electric potentials generated by active axons at each recording electrode. We then systematically varied technical factors (e.g., pulse width, waveform shape, interphase interval (IPI), stimulation configuration, and stimulation frequency) and anatomical parameters (e.g., dorsal cerebrospinal fluid (dCSF) thickness, electrode impedance, electrode positioning relative to vertebrae).

Our results demonstrate that both anatomical and technical factors may significantly alter ECAP amplitude, morphology, and timing. We demonstrate that the choice of recording electrodes can be leveraged to optimize ECAP signals. Additionally, stimulation parameters introduced tradeoffs between axon selectivity, activation thresholds, and ECAP morphology and timing. Importantly, we found that similar ECAP amplitudes can result from distinct neural activation patterns, highlighting the need to interpret ECAPs within their full anatomical and technical context. In some cases, complex ECAP morphologies reflected the spatiotemporal spread of neural recruitment, which may be used to inform our mechanistic understanding of SCS and guide clinical interpretations. These findings are intended to provide engineers and clinicians with relevant insights into the effects of these parameters, supporting the continued development of ECAPs as real-time feedback signals in SCS systems and offer a mechanistic foundation for optimizing stimulation strategies, hardware design, and programming.

## 2. Materials and methods

In this study we utilized a computational model of SCS to systematically characterize the effects of stimulation parameters, recording configurations, and anatomical variations on dorsal column activation and SCS-induced ECAPs [11,15]. We adopted a hybrid modeling approach, combining FEM models of the lower thoracic spine with multicompartment axon models to simulate the neural response to SCS.

### 2.1. Computational model of SCS-induced ECAPs

Our FEM model consisted of the lower thoracic spinal cord, including white and gray matter, cerebrospinal fluid (CSF), dura, epidural tissue, vertebral bone, intervertebral discs, and a surrounding bulk tissue domain (**Figure 1**) [11,15,29]. We positioned an eight-electrode, 75-mm long cylindrical SCS electrode array within the dorsal epidural tissue along the anatomical midline surrounded by a 0.3-mm thick encapsulation layer [15,29,35]. Each electrode was 3-mm long, with a diameter of 1.3 mm and a 4-mm edge-to-edge spacing between electrodes [11]. We defined the electrodes as E0 (most rostral) to E7 (most caudal).

**Figure 1:**
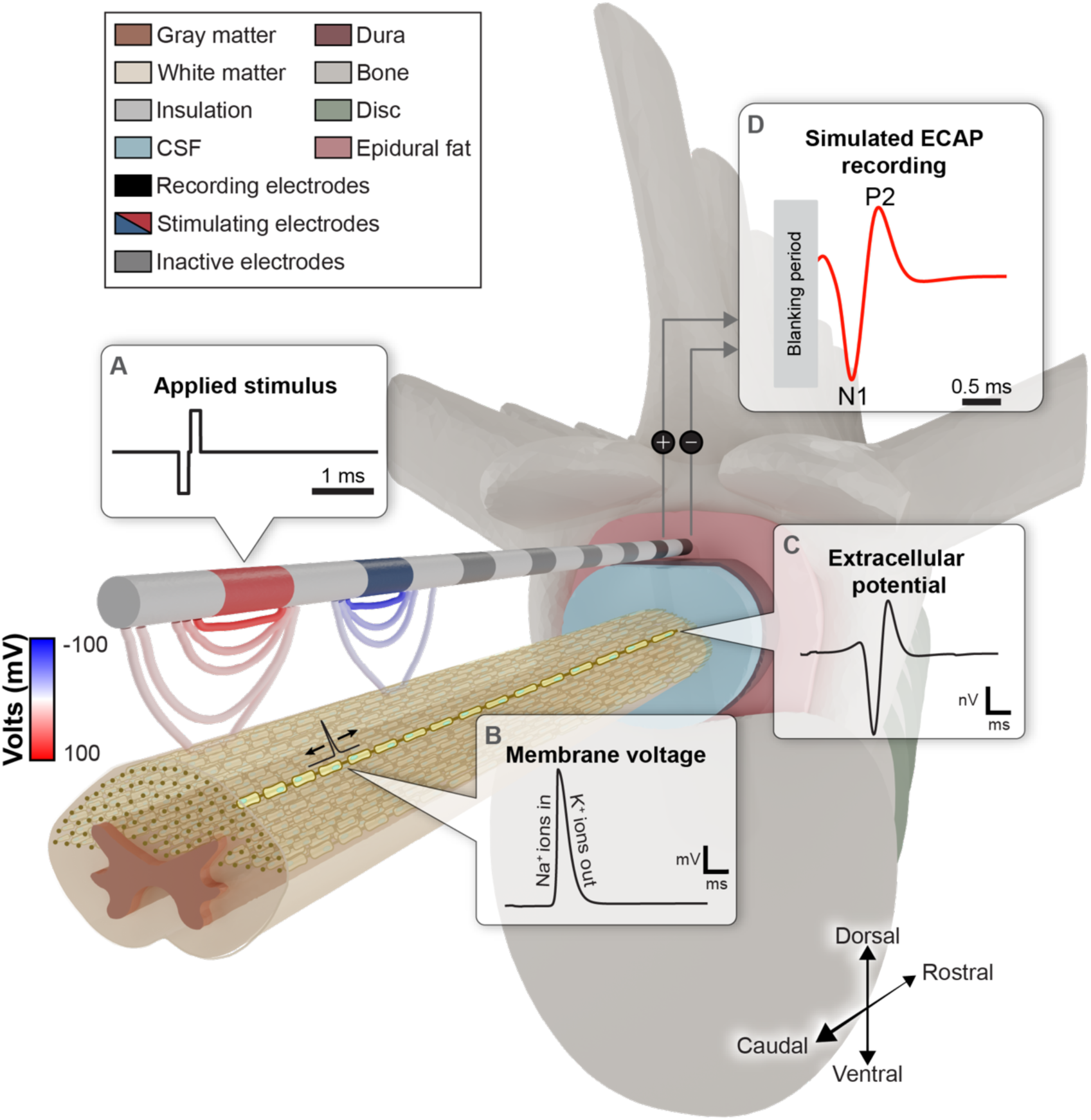
Computational modeling of evoked compound action potentials (ECAPs) generated during SCS. We utilized a previously published model of the lower thoracic spinal cord, surrounding anatomy, and an SCS electrode array to simulate ECAP recordings [11,15,29]. We modeled axons with multicompartment cable models distributed throughout the dorsal half of the spinal cord. Axial positions of the axons are represented as brown circles on the caudal end of the spinal cord, with all fibers running in parallel trajectories along the rostrocaudal axis. One exemplary axon is highlighted in yellow. (A) To simulate the neural response to SCS, we first calculated the extracellular potentials generated by the applied stimulation pulse. (B) We applied these potentials to each modeled axon to simulate membrane voltage responses. (C) We then calculated the extracellular potentials generated at the recording electrodes for each simulated axon. (D) Finally, we generated ECAPs by summing the contributions of all active axons at the recording electrodes, scaled according to the axon diameter using physiologic densities [15,48].

We discretized our model in 3-matic (Materialise, NV, Belgium) to generate a tetrahedral mesh. We then exported the mesh into COMSOL Multiphysics (COMSOL Inc., USA) and assigned tissue conductivities in accordance with previous literature [15]. We modeled the electrode shaft as a perfect insulator and grounded the outer boundary of the general tissue domain. To calculate electrostatic solutions, we applied a unit stimulus (i.e., 1 A) to a single electrode while setting equipotential boundary conditions to all inactive electrodes. We then used the conjugate gradient method to solve for the extracellular electric potentials resulting from stimulation applied at each electrode. Finally, we superimposed individual potential field solutions to calculate the overall potential fields generated by each stimulation configuration.

We simulated axons using a previously published multicompartment sensory axon model [15,36–39]. Using Lloyd’s algorithm, we virtually distributed 100 axons in the dorsal half of the white matter of the spinal cord [29,40]. We used these 100 axial positions within the spinal cord to simulate axons with diameters from 6.0 to 15.9 µm in 0.1 µm increments (i.e., 100 axons per fiber diameter), for a total of 10,000 axons (**Figure 1**) [11]. We performed all axon simulations using the NEURON package in Python 3.9.4 [41,42].

We used the theorem of reciprocity to generate model ECAP recordings (**Figure 1**) [15,43–47]. Specifically, we first applied time-dependent extracellular potentials generated by the stimulating electrodes to each compartment of every modeled axon to the determine the axon’s response to the applied stimulus (**Figure 1A-B**). Next, we calculated the electric potentials at the recording electrodes using the unit stimulus solutions from each recording electrode as transfer functions. These transfer functions served to scale the transmembrane currents from each axonal compartment to determine the resulting single-axon recordings (**Figure 1C**). We calculated the total ECAP recording by summing the recordings from all active axons, scaled by physiologic densities [15,29,48]. To save computational space and time, we excluded voltages generated by inactive axons as their inclusion resulted in minimal differences to model ECAPs (data not shown) [11]. Consistent with clinical recordings of ECAPs [11,12,23,24,49–51], we calculated the final ECAP recording using a differential bipolar recording configuration (**Figure 1D**).

For each ECAP recording, we blanked the stimulation artifact from the onset of the stimulus until 200 µs following the end of the stimulus pulse (**Figure 1D**) [23]. We calculated the ECAP amplitude as the difference between the maximum (i.e., P2) and minimum peak (i.e., N1) voltages within a time window following the blanking period up to 3.0 ms after the onset of the stimulus (**Figure 1D**). We defined the model ECAP threshold (ET) as the lowest stimulation amplitude at which the ECAP amplitude exceeded 4 µV [52,53]. We used ET as a proxy for the perception threshold (PT) [49], and defined the model discomfort threshold (DT) as 1.4*ET [30,31,54–56]. Our primary analysis focused on ECAP characteristics within a paresthesia window from ET to DT.

Growth curves are a well-established method for characterizing ECAPs as a function of stimulation amplitude, providing quantitative insight into changes in neural recruitment and dynamics [11,23,49,57]. Furthermore, growth curves may be utilized to optimize closed-loop SCS dosing [53,58,59]. Therefore, we generated model growth curves for each parameter set by plotting ECAP amplitudes at each stimulation amplitude from 0.1 mA to DT in 0.1 mA increments.

Closed-loop SCS systems aim to achieve consistent neural activation using ECAP amplitudes [9,10,59,60]. However, ECAP amplitude alone does not always accurately reflect similar levels of neural recruitment. For example, changes in patient posture or variations in dCSF thickness may significantly alter the underlying neural activation without causing corresponding changes in ECAP amplitude [11]. Similarly, adjustments in stimulation parameters, such as stimulation configuration, can modulate the spatiotemporal patterns of axon recruitment, resulting in ECAPs with differing morphologies [12,17,50]. To better understand the relationship between ECAPs and underlying neural activation, we generated curves of activated axon counts across stimulation amplitudes and ECAP amplitudes. In conjunction with growth curves, we used these curves to evaluate whether similar stimulation or ECAP amplitudes equated to similar levels of neural activation. Finally, for a subset of parameters, we extended our analysis of ECAP composition by examining the spatiotemporal properties of axonal activation and decomposing the total ECAP according to axon diameter or location of activation. Consistent with the scaling for our ECAP calculations, we scaled the number of activated fibers according to physiologic densities [43–45,48].

### 2.2. Model parameter evaluation

Contemporary SCS devices have enabled extensive customization of both stimulation and recording parameters, including stimulation waveform shape, IPI, pulse width, stimulation frequency, and electrode configurations. Each of these parameters may significantly influence the neural response to stimulation and alter the characteristics of recorded ECAPs. Anatomical variability (e.g., changes in dCSF thickness and electrode position relative to vertebrae) may further modulate dorsal column activation and complicate the interpretation of ECAPs [11–15,23,24,49]. Therefore, our objective was to systematically characterize the effects of individual parameter variations on SCS-induced ECAPs and the underlying neural responses.

To accomplish this, we defined a set of computational modeling experiments in which we systematically swept through anatomical, stimulation, and recording parameter spaces. We established a base model with a 3-mm dCSF thickness and a bipolar stimulation configuration (E6+/E7-), which is frequently used in studies of SCS-induced ECAPs [12,13,16,50,61]. We applied stimulation using a 50 Hz, symmetric biphasic rectangular pulse with a 150-µs pulse width and 30-µs IPI (**Figure 1A**). We selected 50 Hz as the stimulation frequency to approximate the frequency used in conventional SCS and studies of ECAPs recorded during SCS [12,15,23,24,49,62–64]. We set the base model conductivity of the encapsulation layer to 0.089 S/m to align the model median bipolar electrode impedance to the median bipolar impedance in a clinical ECAP dataset (911 Ω) [11,35,65].

Full details of the tested parameters and values are provided in **Table 1**. We evaluated the effects of varying each parameter individually, with all other settings held constant as defined in the base model.

**Table 1:** Parameters tested in the computational modeling analyses. Bolded values indicate the base model. When evaluating the model response to each parameter, all other parameters were held constant at the value used in the base model.

| Tested parameter |  |  |
| --- | --- | --- |
| Stimulation configuration |  | Monopolar (E6-)<br>**Bipolar (E6+/E7-)**<br>Wide bipolar (E5+/E7-)<br>Guarded cathode (E5+/E6-/E7+) |
| Stimulation waveform | Morphology | **Biphasic rectangular**<br>Triphasic rectangular |
| | Pulse width (μs) | 90, 120, **150**, 180, 210, 240, 270, 300 |
|  | Triphasic 1<sup>st</sup>/3<sup>rd</sup> phase contribution (%/%) | **0.0/100.0**, 16.7/83.3, 33.3/66.7, 50.0/50.0, 66.7/33.3, 83.3/16.7, 100.0/0.0 |
| | Inter-phase interval (μs) | 0, **30**, 60, 90, 120 |
|  | Pulse frequency (Hz) | **50**, 100, 150, 500, 1000 |
| Recording configuration |  | **E1-E0**<br>E2-E0<br>E3-E0 |
| Anatomical parameters | Dorsal cerebrospinal fluid thickness (mm) | 1, 2, **3**, 4, 5 |

### 2.3. Recording configurations

We used our computational model to analyze several differential recording pairs (E1-E0, E2-E0, and E3-E0). To investigate the underlying causes of these changes, we examined the monopolar recordings (E0, E1, E2, and E3) and assessed how each contributed to components of the differential recordings.

### 2.4. Anatomical factors

Variability in the thickness of the dCSF layer, both on a patient-specific basis and in response to physiological factors, such as posture and respiration, present a significant challenge for SCS systems. Fluctuations in dCSF thickness alter the distance between the implanted electrode arrays and the spinal cord, which may result in over- or under-stimulation [11,15,66,67]. These variations are critical as they can modulate the neural response to SCS, including perception thresholds and ECAPs, ultimately affecting the efficacy of SCS therapies [11,15,68,69]. To evaluate the impact of dCSF thickness on axon recruitment and ECAPs, we generated computational models with dCSF thicknesses ranging from 1 to 5 mm in 1-mm increments, which spanned previously reported ranges [66,69].

### 2.5. Stimulation waveform parameters

The selection of stimulation waveform parameters in SCS systems varies considerably, reflecting a wide range of clinical practices and device capabilities. To optimize both open-loop and closed-loop SCS, it is essential that we understand how each parameter influences the underlying neural response and how these effects are reflected in SCS-induced ECAPs.

Conventional tonic SCS employs charge-balanced waveforms with a range of pulse widths [11,12,23,49,70,71]. Building on prior work [11], we evaluated the effects of pulse width variations from 90 to 300 µs in 30-µs increments. Additionally, studies using monopolar stimulation have found that introducing an interphase gap can reduce biphasic stimulation thresholds comparable with monophasic stimulation [72–74], however, other evidence suggests that stimulating with the more commonly used bipolar stimulation configuration may raise thresholds as IPIs are increased [75]. To our knowledge, the influence of IPIs on ECAPs generated during SCS has not been well characterized. Therefore, we further employed our model to examine how varying IPIs from 0 to 120 µs in 30-µs intervals influenced neural recruitment and ECAP characteristics.

In addition to the temporal parameters of the applied stimulus, waveform morphology and polarity can influence neural responses to SCS. While symmetric, biphasic rectangular pulses are widely used for their charge balance and efficiency, alternative waveforms have been proposed to reduce stimulation artifact, modulate recruitment patterns, or optimize ECAP measurements [13,14,17,20,50,76]. Additionally, asymmetric waveforms have been proposed to reduce stimulation artifact in ECAP recordings by shifting activation toward the end of the stimulation pulse [13,14,17]. However, ECAP morphology can vary with the duration, polarity, and timing of the stimulation phases. We hypothesized that asymmetric triphasic waveforms may modulate the spatiotemporal dynamics of fiber activation and thereby alter ECAP morphologies. Therefore, we evaluated the effects of waveform morphology by comparing biphasic-rectangular and triphasic-rectangular waveforms. We used our computational model to vary the relative durations of the first and third phases of the rectangular triphasic waveform while holding the second, activating phase constant at 150 µs. To maintain charge balance, the total duration of the first and third phases summed to 150 µs. We defined five triphasic waveforms with first-phase contributions of 16.7% (∼25 µs) to 83.3% (∼125 µs) in 16.7% increments. We included two biphasic waveforms in our analysis: a cathodic-leading biphasic pulse (base model, representing a 0% first-phase contribution in the triphasic waveform), and an anodic-leading biphasic pulse (representing a 100% first-phase contribution).

Current closed-loop SCS systems primarily deliver stimulation using bipolar or tripolar configurations, however, alternative stimulation configurations may offer advantages in terms of selective recruitment and efficiency [12,49,50,76]. Given that modern SCS systems support a wide range of programmable configurations, it is essential to understand how neural responses and ECAPs vary across configurations. This understanding is also clinically relevant, as patients may respond differently or have preferences for certain configurations [77,78]. Therefore, we evaluated four stimulation configurations to assess their effects on axonal recruitment and ECAP characteristics during SCS. These included two configurations commonly used in clinical studies of SCS-induced ECAPs, bipolar and guarded-cathode [11,12,23,27,49–51], as well as monopolar and wide-bipolar configurations.

Recent work has highlighted frequency-dependent characteristics of ECAPs and underlying neural activation during SCS [21,27]. However, studying these effects experimentally, particularly at higher frequencies, is challenging due to sources of artifact and biological noise which obscure the ECAP component of the recordings. Computational models offer a valuable alternative by enabling detailed analysis of the neural-specific components of ECAPs and the ability to isolate the neural-specific components of the recorded signal and individual fiber responses. Therefore, we used our base model to examine how ECAPs and the underlying neural firing evolve with increasing stimulation frequency. We performed simulations at five pulse frequencies: 50, 100, 150, 500, and 1000 Hz. We applied stimulation for 100 ms at amplitudes ranging from 0.1 mA up to DT for each stimulation frequency.

We generated ECAP growth curves for individual stimulus pulses using the difference between the maximum (i.e., P2) and minimum peak (i.e., N1) voltages within a time window following the blanking period up to the start of the subsequent stimulus pulse. We then generated averaged growth curves using the average ECAP amplitudes across all applied pulses for each stimulation frequency. We defined ET as the lowest stimulation amplitude at which the ECAP amplitude exceeded 4 µV in the average growth curve, calculated across applied pulses for each respective stimulation frequency. All stimulation pulses were identical across frequencies and consisted of symmetric biphasic waveforms with a 150-µs pulse width and 30-µs IPI.

## 3. Results

### 3.1. Recording configurations

To examine the effect of recording configurations on ECAPs, we tested three bipolar recording configurations: E1-E0, E2-E0, and E3-E0. Recording configurations E2-E0 and E3-E0 yielded larger ECAP amplitudes and shorter peak latencies than E1-E0 (**Figure 2B-C**). The increase in ECAP amplitudes reduced ET from 3.8 mA for E1-E0 to 3.5 mA for both E2-E0 and E3-E0 (**Figure 2B-C**). DTs decreased in a similar manner from 5.3 mA with E1-E0 to 4.9 mA for E2-E0 and E3-E0. Recording on E3-E0 consistently required the fewest number of active axons to generate a given ECAP amplitude (**Figure 2D**).

**Figure 2:**
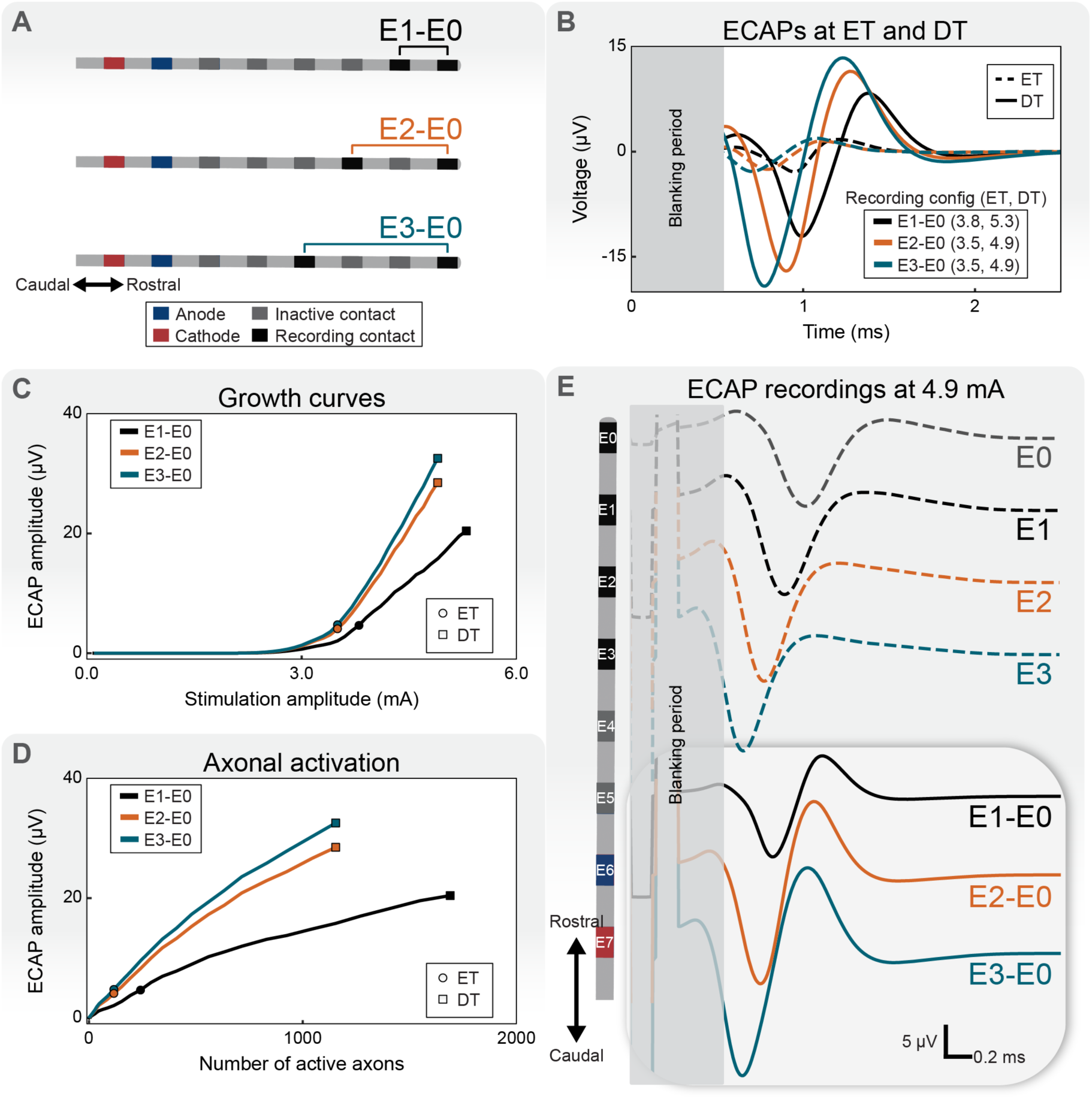
Influence of recording configuration on evoked compound action potentials (ECAPs). (A) We simulated dorsal column responses and ECAP recordings for three differential bipolar recording configurations: E1-E0 (black), E2-E0 (orange), and E3-E0 (blue). (B) ECAPs at ECAP threshold (ET) and discomfort threshold (DT). ETs and DTs are shown in mA. (C) Growth curves for each recording configuration. (D) ECAP amplitude as a function of axonal activation. (E) To further investigate differences across recording configurations, we compared monopolar recordings (top) to differential bipolar recordings (bottom) in response to stimulation at 4.9 mA.

Next we analyzed monopolar recordings from each recording electrode (E0, E1, E2, and E3) (**Figure 2E**). We applied stimulation at 4.9 mA, which corresponded to DT for the E2-E0 and E3-E0 recording configurations. At 4.9 mA, recording from E3 produced an ECAP amplitude of 20.2 µV. Recording 21 mm rostral at E0 resulted in an ∼27% reduction in the ECAP amplitude, to 14.8 µV. These differences highlight the effects of proximity of the recording site to the stimulation site on both ECAP amplitude and temporal dispersion.

In differential recordings, we observed the largest ECAP amplitudes when inter-electrode spacing aligned opposing ECAP peaks in time across electrodes (**Figure 2E**). Specifically, N1 was maximized in the differential recording when the peak of the negative phase reached E3, the electrode proximal to the stimulation electrodes, while the distal electrode E0 captured the leading positive peak of the propagating waveform. Additionally, the P2 peak was maximized when the trailing positive peak reached E3 concurrently with the negative peak at E0. This temporal offset, combined with the proximity of E3 to the stimulation site, maximized the differential signal thereby increasing the ECAP amplitude (**Figure 2E**).

### 3.2. Anatomical factors: dCSF thickness, rostrocaudal electrode position, and electrode impedance

dCSF thickness had a substantial impact on neural recruitment and ECAP characteristics (**Figure 3**). ETs and DTs were highly sensitive to changes in dCSF thickness. Shifting the dCSF thickness by 4 mm (from 1 to 5 mm) resulted in an approximately 640% increase in both ET (1.2 to 8.9 mA) and DT (1.7 to 12.5 mA) (**Figure 3B-C**). Larger dCSF thicknesses shifted growth curves toward higher stimulation amplitudes and reduced the slope of their linear portions (i.e., between ET and DT) as the distance between the spinal cord and implanted electrode array increased (**Figure 3C**).

**Figure 3:**
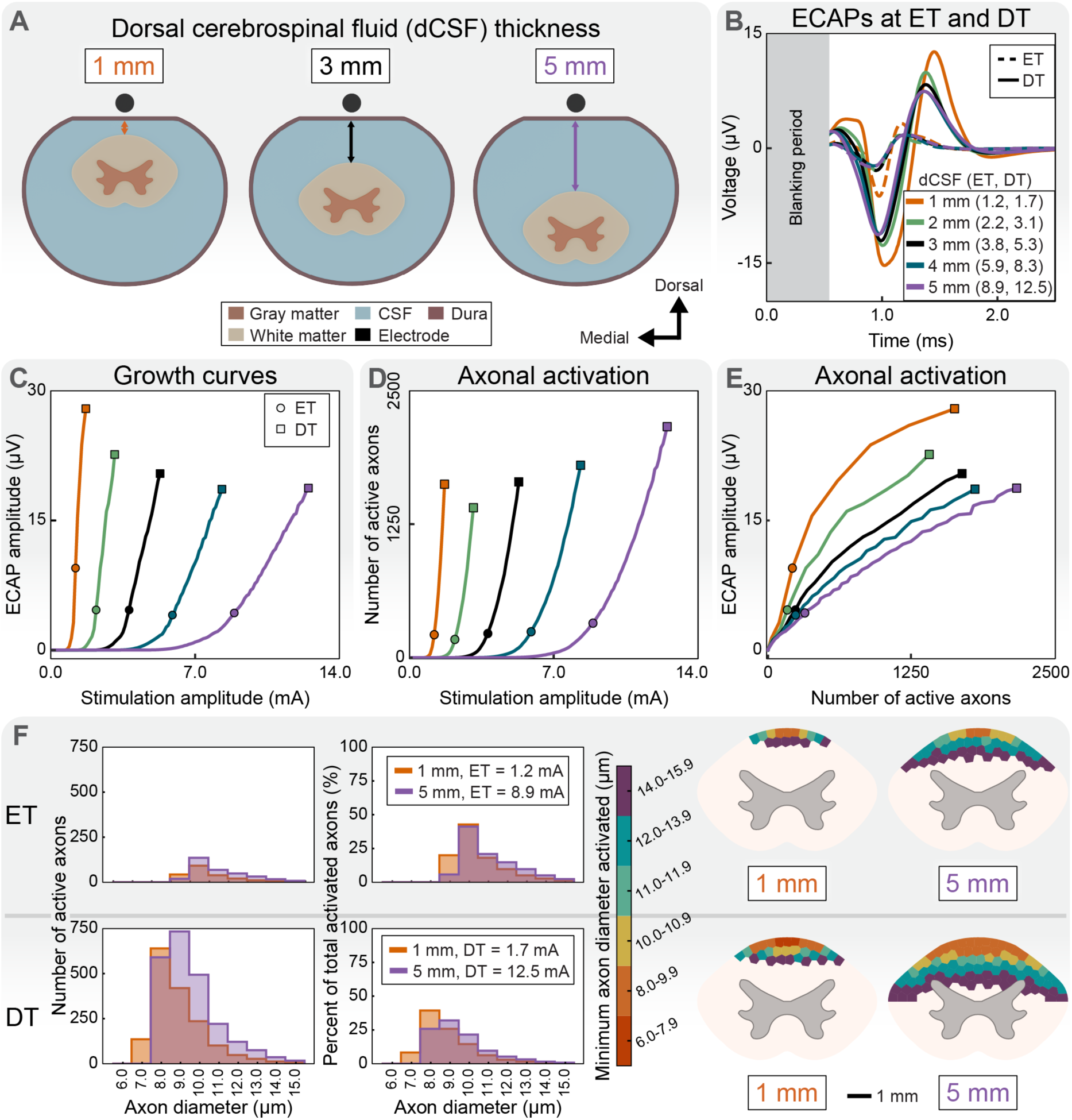
Effects of dCSF thickness on evoked compound action potentials (ECAPs) and axonal activation. (A) We computed neural responses and model ECAPs for dCSF thicknesses ranging from 1 to 5 mm in 1-mm increments. (B) ECAPs at ECAP threshold (ET) and discomfort threshold (DT). ETs and DTs are shown in mA. (C) Growth curves for each dCSF thickness. Note that the ET for the 1-mm dCSF model is larger than 4 µV due to the granularity of the stimulation current step size (0.1 mA). (D) Axonal activation as a function of stimulation amplitude. (E) ECAP amplitude as a function of axonal activation. (F) To investigate whether dCSF thickness influenced the recruitment of different axon populations, we compared the recruitment profiles of the 1-mm and 5-mm dCSF models at ET (top row) and DT (bottom row). Histograms of activated axons by diameter (left) showed that the 1-mm dCSF model activated a larger proportion of small-diameter axons at both ET and DT. Cross-sectional maps of the minimum activated axon diameter (right) indicated this difference was confined to the superficial dorsal columns, whereas the 5-mm dCSF model activated a broader region as well as more large-diameter axons.

The number of active axons at ET and DT also increased with dCSF thickness (**Figure 3D-E**). At ET, the number of active axons ranged from 170 (at 2.2 mA; 2-mm dCSF model) to 322 axons (at 8.9 mA; 5-mm dCSF model). The 1-mm dCSF model activated more axons at ET (214 fibers) than the 2-mm model due to steeper slope of the growth curve and the stimulation amplitude step size employed in this study (0.1 mA – a standard step size in clinical SCS systems). Specifically, in the 1-mm dCSF model, the ECAP amplitude increased rapidly from 3.2 µV at 1.1 mA to 9.5 µV at 1.2 mA, which was the largest ECAP amplitude observed at ET across all dCSF thicknesses (**Figure 3C**). Across all dCSF thicknesses at DT, active axon counts ranged from 1405 (at 3.1 mA, 2-mm dCSF model) to 2164 axons (at 12.5 mA, 5-mm dCSF model) (**Figure 3D-E**). The 1-mm dCSF model consistently required fewer active axons to generate equivalent ECAP amplitudes than larger dCSF thicknesses (**Figure 3E**).

To characterize neural recruitment, we generated histograms of active axon counts and cross-sectional maps of the spinal cord with the minimum axon diameters active per area for the 1-mm and 5-mm dCSF models (**Figure 3F**). Although the 5-mm dCSF model activated a greater total number of axons at both ET and DT, the 1-mm dCSF model preferentially recruited more smaller-diameter axons in the superficial dorsal columns near the electrodes.

### 3.3. Stimulation waveform parameters

Increasing the pulse width from 90 to 300 µs reduced both ETs and DTs by approximately 60% and delayed peak timings by 20-29% (**Figure 4B**). Specifically, at ET, the N1 and P2 peaks were delayed by 0.25 ms and 0.22 ms, respectively. At DT, the N1 peak was delayed by 0.22 ms and the P2 peak was delayed by 0.30 ms when comparing stimulation with pulse widths of 90 versus 300 µs. Longer pulses widths also broadened the ECAP waveform and shifted growth curves toward lower stimulation amplitudes (**Figure 4B-C**). These effects were most prominent for shorter pulse widths (i.e., from 90 to 150 µs). In addition, increasing the pulse width consistently lowered activation thresholds and increased the number of recruited axons per stimulation amplitude (**Figure 4D**). ECAP amplitudes, particularly for pulse widths shorter than 180 µs, reflected similar levels of neural recruitment (**Figure 4E**).

**Figure 4:**
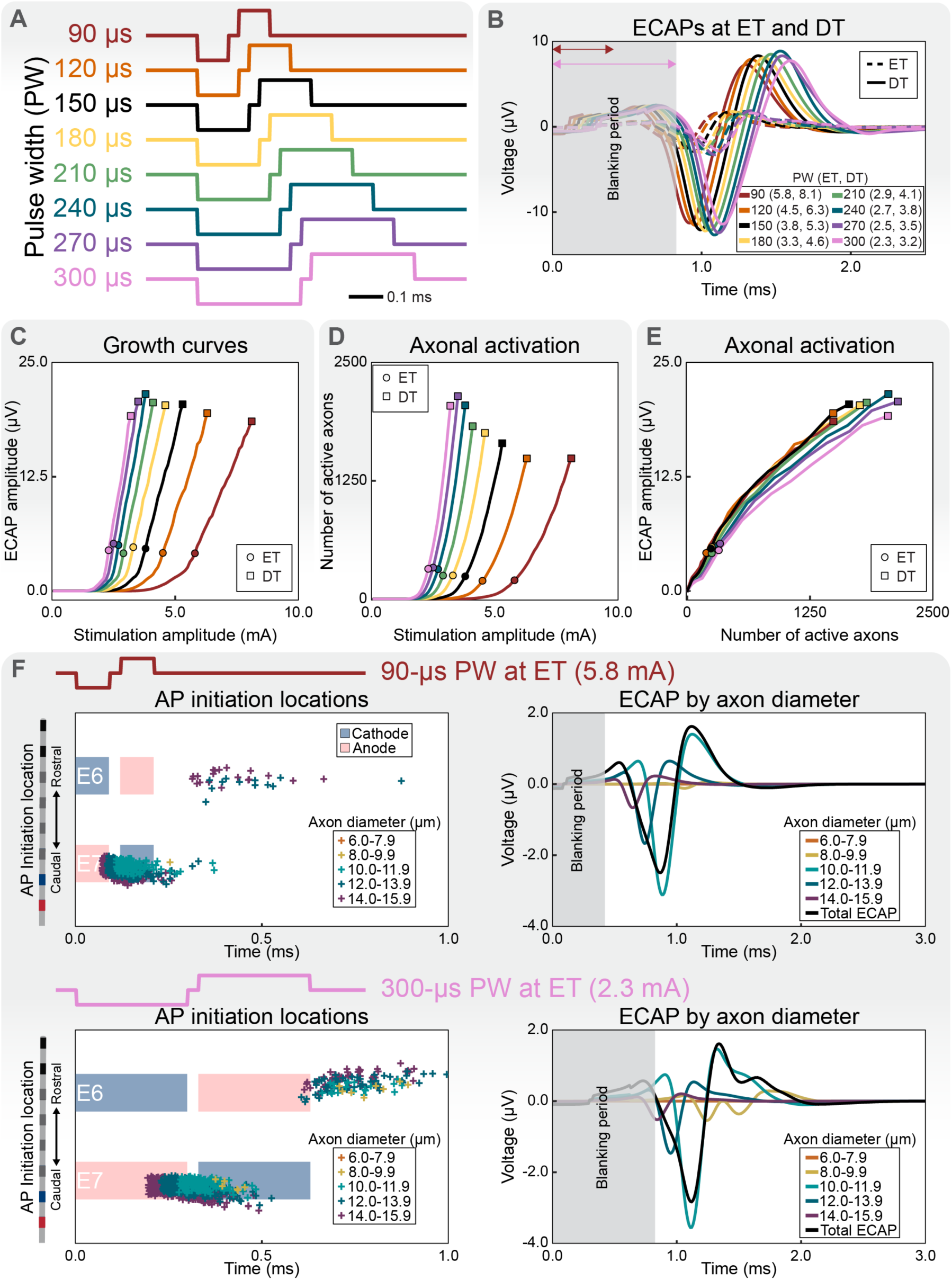
Effects of stimulation pulse width (PW) on evoked compound action potentials (ECAPs) and axonal activation. (A) We simulated neural responses and ECAPs for PWs ranging from 90 to 300 µs in 30-µs increments. (B) ECAPs at ECAP threshold (ET) and discomfort threshold (DT) for each PW. ETs and DTs are shown in mA. For each PW, we blanked the signal from the start of stimulation until 200 µs following the end of the stimulus. The blanking period shown in gray corresponds to the time window for the 300-µs PW (pink arrow). The blanking period is also overlaid with a red arrow to indicate the blanking time for the 90-µs PW. (C) ECAP growth curves for each PW. (D) Axonal activation as a function of stimulation amplitude. (E) ECAP amplitude as a function of axonal activation. (F) We calculated the action potential (AP) initiation time and location for each active axon at ET for the 90-µs (top, left column) and 300-µs PW (bottom, left column). Shaded regions indicate each phase of the stimulation waveform, with the cathode and anode shown in red and blue, respectively. We then separated the total ECAP according to signal contributions from each axon-diameter group (right column).

To further examine the factors underlying the observed differences in ECAPs, we analyzed the spatial and temporal patterns of neural activation at ET for pulse widths of 90 and 300 µs (**Figure 4F**). Specifically, we decomposed the ECAPs according to the timing and location of action potential initiation (**Figure 4F, left**). At ET for the 90-µs pulse width (5.8 mA), the ECAP was predominantly composed of activation that occurred near the cathode (E7) during the first phase of stimulation (**Figure 4F, top left**). Axons with diameters between 10.0 and 11.9 µm had the largest contribution to the total recorded ECAP (**Figure 4F, top right**). In contrast, at ET for the 300-µs pulse width (2.3 mA), 60% more active axons were required to obtain an ECAP amplitude above the threshold for ET (200 versus 320 axons for pulse widths of 90 and 300 µs, respectively). Similar to the 90-µs pulse width, stimulation with a pulse width of 300 µs resulted primarily in activation near E7 during the first phase of stimulation (**Figure 4F, bottom**). However, in addition to the 238 axons activated during the first phase at ET, stimulation with a pulse width of 300 µs activated an additional 82 axons during the second phase of stimulation. This additional recruitment of smaller-diameter axons introduced an additional peak following P2 in the total ECAP (**Figure 4F, bottom right**).

Next, we considered the role of pulse timing by varying IPIs from 0 to 120 µs in 30-µs intervals (**Figure 5A**). We found that increasing the IPI from 0 to 120 µs decreased ET and DT by ∼10%, with the largest difference in thresholds between IPIs of 0 and 30 µs (**Figure 5B**). Beyond 30 µs, ETs decreased by 0.1 mA for every 30-µs increase in IPI until plateauing for IPIs longer than 90 µs (**Figure 5B-C**). As IPI increased, growth curves shifted toward lower stimulation amplitudes (**Figure 5C**). Again, this effect was most pronounced at shorter IPIs (i.e., 0 and 30 µs) and diminished at longer IPIs (i.e., 90 and 120 µs). Although the 120-µs IPI recruited the greatest number of axons per stimulation amplitude, activation was similar for IPIs above 60-µs (**Figure 5D**). Interestingly, while the 120-µs IPI produced the lowest ET, DT, and the largest ECAP amplitudes, it paradoxically required the largest number of axons to achieve a given ECAP amplitude (**Figure 5E**). Upon further analysis, we found that increasing the IPI lowered activation thresholds and, at low stimulation amplitudes, increased the proportion of axons activated during the second phase of stimulation (**Figure 5F**). When applying stimulation with a bipolar stimulation configuration (E6+/E7-) using a 120-µs IPI, the first phase of stimulation activated axons near E7, and subsequent activation occurred near E6 during the second phase of stimulation (**Figure 5F, bottom**). This temporal and spatial offset led to partial cancellation in the total recorded ECAP. As a result, ∼56% more axons were recruited at ET for the 120-µs IPI (3.6 mA; 311 active axons) compared to the 0-µs IPI (4.0 mA, 200 active axons). This effect diminished at higher stimulation amplitudes, as activation predominantly occurred during the first phase of stimulation.

**Figure 5:**
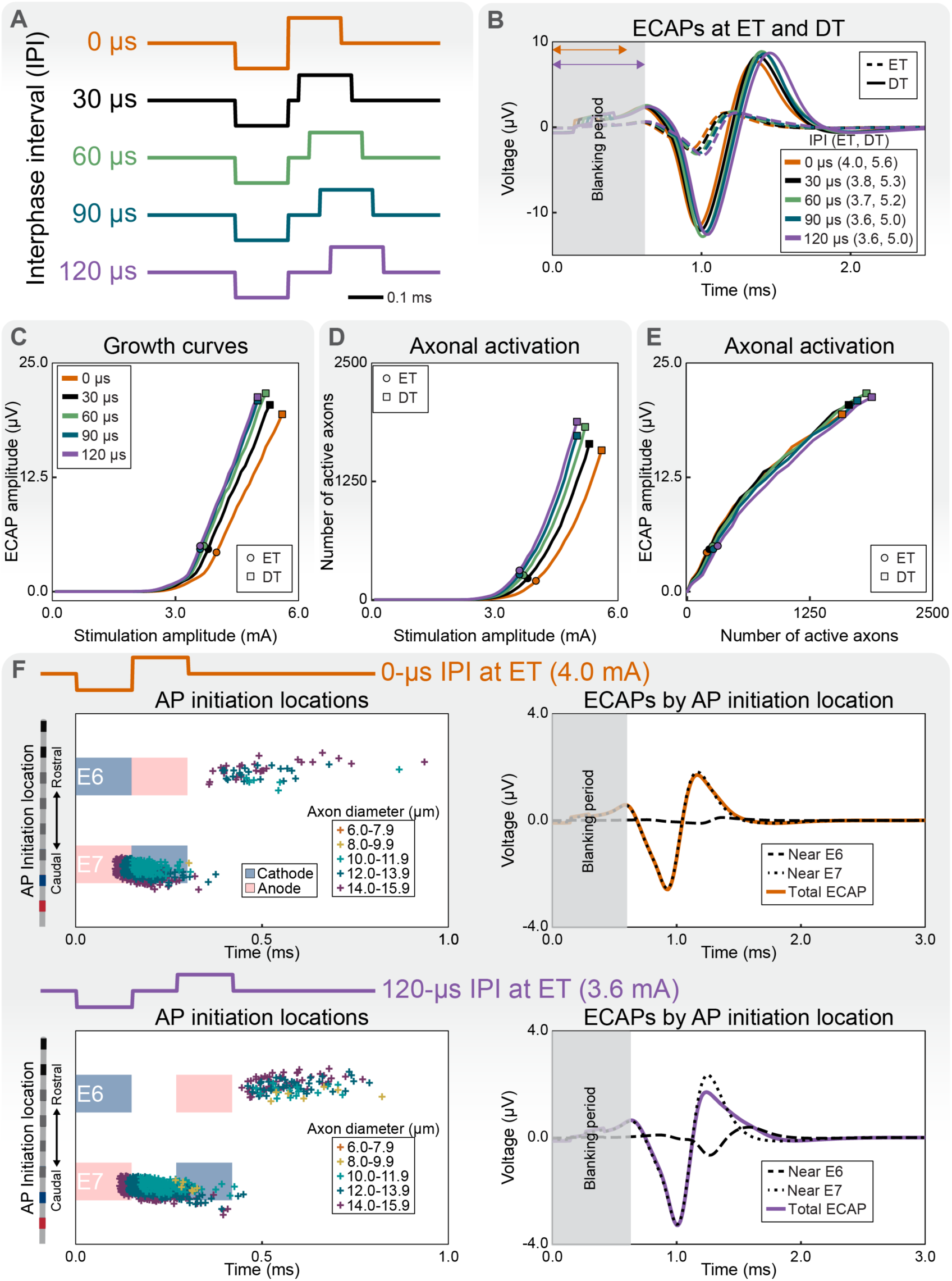
Effects of interphase interval (IPI) on evoked compound action potentials (ECAPs) and axonal activation. (A) We simulated neural responses and ECAPs for IPIs from 0 to 120 µs in 30-µs intervals for a fixed 150-µs pulse width. (B) ECAPs at ECAP threshold (ET) and discomfort threshold (DT). ETs and DTs are shown in mA. For each IPI, we blanked the signal from the start of stimulation until 200 µs following the end of the stimulus. The blanking period shown in gray corresponds to the time window for the 120-µs IPI (purple arrow). The blanking period is overlaid with an orange arrow to indicate the blanking time for the 0-µs IPI. (C) Growth curves for each IPI. (D) Axonal activation as a function of stimulation amplitude. (E) ECAP amplitude as a function of axonal activation. (F) The action potential (AP) initiation time and location for each active axon at ET for the 0- and 120-µs IPI (left). Shaded regions indicate each phase of the stimulation waveform, with the cathode and anode shown in red and blue, respectively. We decomposed the resulting ECAPs according to the location of activation (right).

Next, we evaluated the effects of waveform morphology by comparing biphasic rectangular and triphasic rectangular stimulation waveforms. We varied the first-phase contribution of a triphasic waveform relative to the third phase in ∼16.7% (i.e., ∼25 µs) increments, from 0% (i.e., cathodic-leading biphasic) to 100% (i.e., anodic-leading biphasic). The anodic-leading biphasic waveform (i.e., 100% first-phase contribution), equivalent to the E6-/E7+ with the biphasic stimulation in the base model, resulted in the lowest ET and DT (3.7 and 5.2 mA respectively), while the 66.7% triphasic waveform generated the highest thresholds (5.1 and 7.1 mA, respectively) (**Figure 6B-C**). Although stimulation with E6-/E7+ produced the lowest ET and DT, these results do not account for stimulation artifact in clinical recordings, which would increase due to the reduced distance between the activation site (E6) and the recording electrodes. Axonal recruitment was similar across waveforms when comparing ECAP amplitudes, with the exception of the 66.7% triphasic waveform which had reduced ECAP amplitudes with a multiphasic morphology (**Figure 6B-D**). Using model ECAPs, we investigated the underlying neural composition of the ECAPs for the 66.7% triphasic waveform and the cathodic-leading biphasic stimulation (i.e., base model condition) at their respective ETs (**Figure 6F**). Biphasic stimulation produced ECAPs which were primarily composed of activation that occurred near E7, the cathode during the first phase of the stimulation (**Figure 6F, top**). In contrast, the total ECAP generated by the 66.7% triphasic stimulation was composed of activation that occurred near E6 during the first phase of stimulation as well as E7 during the second phase (**Figure 6F, bottom**). The temporal offset combined with the distance between activation sites caused interference in the total recorded ECAP, which lowered the peak-to-peak amplitude. Additionally, the 66.7% triphasic waveform activated 470 axons at ET, nearly double the 241 active axons at ET with the 0% biphasic waveform.

**Figure 6:**
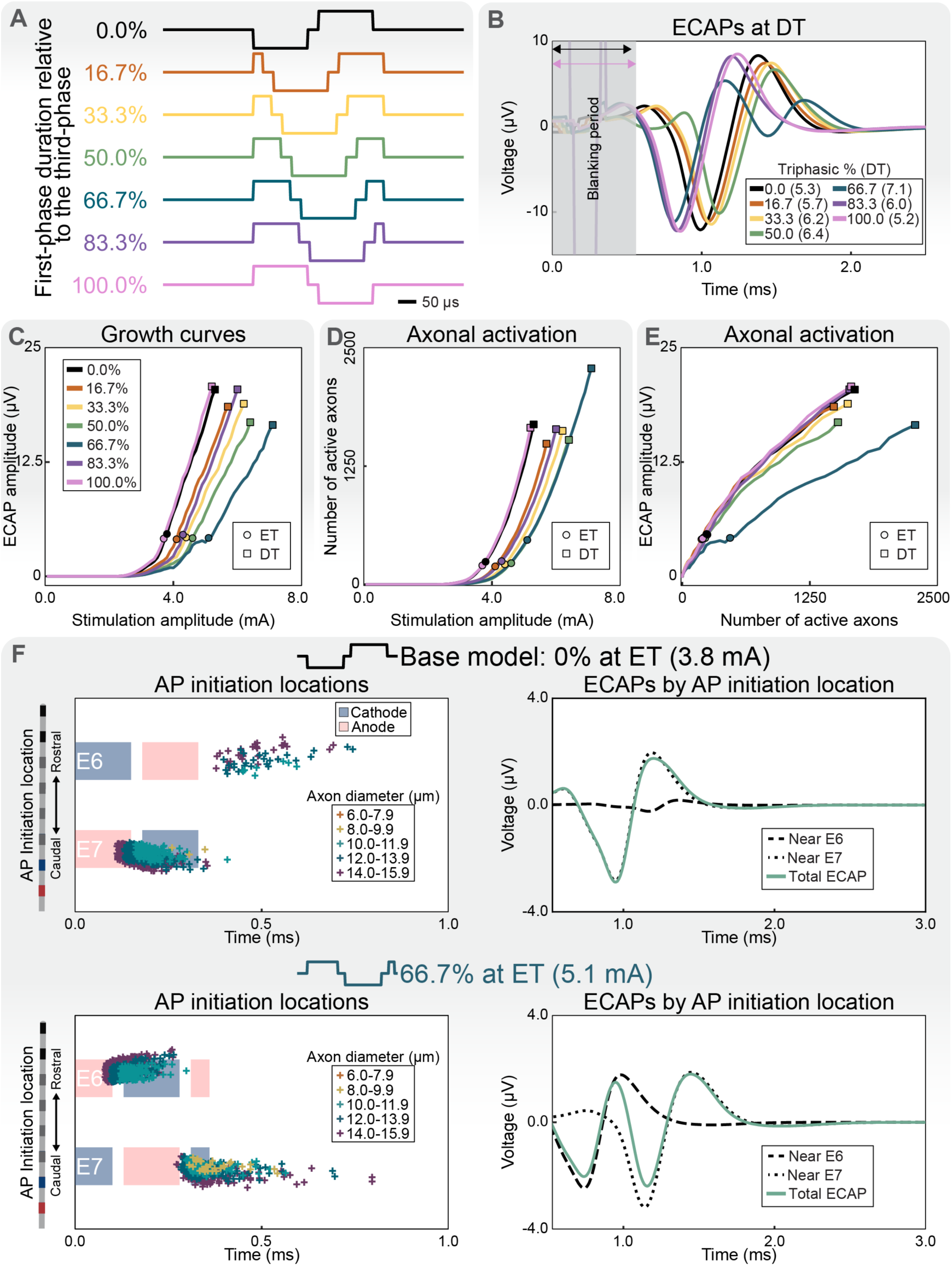
Effects of biphasic stimulation and asymmetric triphasic phase durations on evoked compound action potentials (ECAPs) and axonal activation. (A) We simulated axonal responses and corresponding ECAPs for biphasic and triphasic waveforms. To characterize asymmetric triphasic stimulation waveform morphologies, we systematically varied the relative durations of the first and third phases while holding the middle (second) phase constant at 150 µs. To maintain charge balance, the total duration of the first and third phases was also fixed at 150 µs. We varied the relative contributions of the first and third phases from 0% (cathodic-leading biphasic waveform, i.e., 0-µs first phase) to 100% (anodic-leading biphasic waveform, i.e., 0-µs third phase) in ∼16.7% increments (∼25-µs steps). (B) Modeled ECAPs at discomfort threshold (DT). DTs are reported in mA. ECAPs at ECAP threshold (ET) are provided in **Supplementary Figure 1**. For each waveform, we blanked the signal from the start of stimulation until 200 µs following the end of the stimulus. The blanking period shown in gray corresponds to the time window for the triphasic waveforms (pink arrow). The blanking period is overlaid with a black arrow to indicate the blanking time for the biphasic waveforms. (C) Growth curves across biphasic and triphasic stimulation waveforms. (D) Axonal activation as a function of stimulation amplitude. (E) ECAP amplitude as a function of axonal activation. (F) To investigate the ECAP morphology and growth curve characteristics of the 66.7% triphasic waveform, we compared it to the base biphasic model at each respective ET. For each stimulation waveform, we computed the timing and spatial location of action potential (AP) initiation for all activated axons (left). Shaded regions indicate each phase of the stimulation waveform, with the cathode and anode shown in red and blue, respectively. We then decomposed the resulting ECAPs according to the location of activation (right).

We then compared ECAP characteristics and axonal recruitment across four stimulation configurations: monopolar, bipolar, wide-bipolar, and guarded-cathode. Among the tested stimulation configurations, guarded cathode stimulation generated the lowest ET and DT (3.6 and 5.0 mA, respectively), followed closely by wide bipolar (3.7 and 5.2 mA, respectively) and bipolar stimulation (3.8 and 5.3 mA, respectively) (**Figure 7B-C**). Bipolar, wide bipolar, and guarded cathode recruited similar numbers of axons per stimulation amplitude (**Figure 7D**). Monopolar and bipolar stimulation activated similar numbers of fibers for a given ECAP amplitude, as did guarded cathode and wide bipolar stimulation (**Figure 7E**).

**Figure 7:**
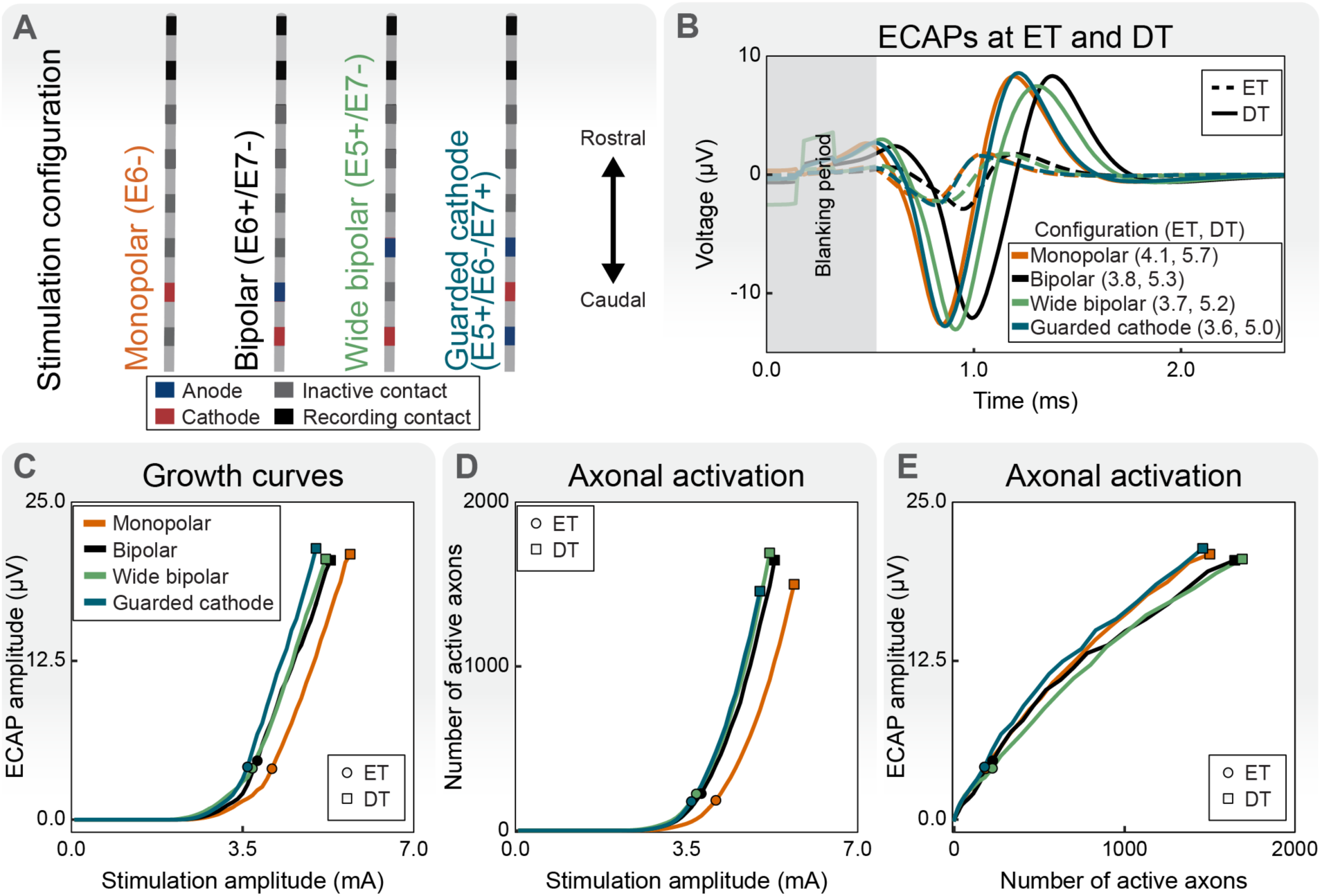
Effects of stimulation configuration on evoked compound action potentials (ECAPs) and axonal activation. (A) We used computational modeling to test the following stimulation configurations: monopolar (E6-), bipolar (E6-/E7-; base model), wide bipolar (E5+/E7-), and guarded cathode (E5+/E6-/E7+). (B) ECAPs at ECAP threshold (ET) and discomfort threshold (DT). ETs and DTs are provided in mA. For each configuration, we blanked the signal from the start of stimulation until 200 µs following the end of the stimulus. (C) ECAP growth curves for each stimulation configuration. (D) Axonal activation as a function of stimulation amplitude. (E) ECAP amplitude as a function of axonal activation.

Finally, we used our base model to examine how ECAPs evolve with increasing stimulation frequency. We performed simulations at five frequencies: 50, 100, 150, 500, and 1000 Hz. We first examined ECAP responses at a fixed stimulation amplitude of 3.8 mA (**Figure 8**), which corresponded to the ET for a pulse frequency of 50 Hz. The temporal activation patterns of fibers differed as the stimulation frequency increased (**Figure 8B-C**). At 50 Hz, once axons began firings they fired in a one-to-one manner with each stimulation pulse, with a median firing rate of 50.0 spikes/second (range: 20 to 50 spikes/second). When stimulating at 500 Hz, axons fired with a median firing rate of 370 spikes/second (range: 30 to 500 spikes/second), while stimulation at 1000 Hz had a median firing rate was 420 spikes/second (range: 20 to 500 spikes/second). This partial entrainment resulted in cyclic modulation of ECAP amplitudes for 500- and 1000-Hz stimulation, where ECAP amplitudes varied across pulses (**Figure 8D**).

**Figure 8.**
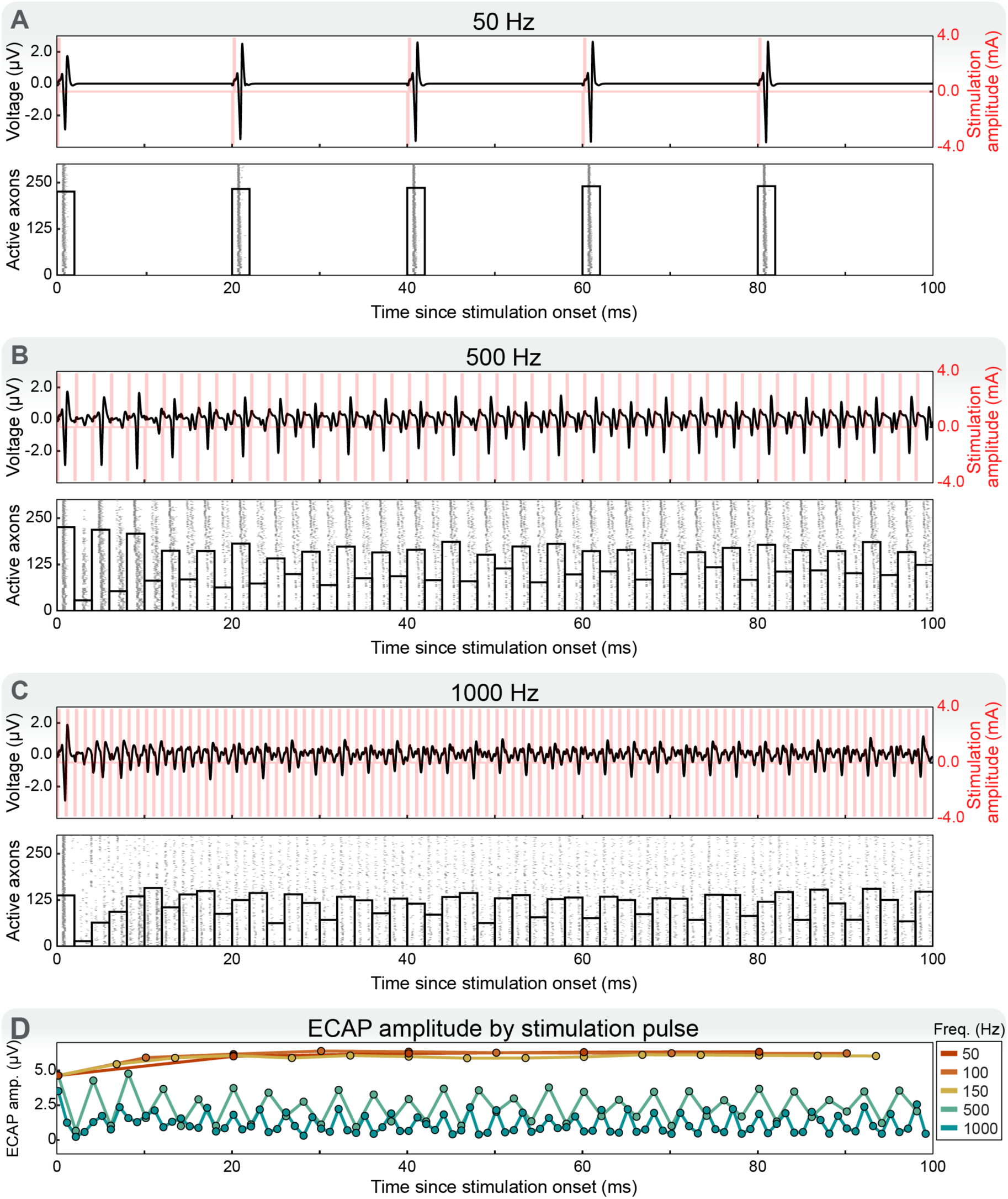
Effect of stimulation frequency on SCS-induced axonal activation profiles and evoked compound action potential (ECAP) recordings. We applied stimulation frequencies of (A) 50 Hz, (B) 500 Hz, and (C) 1000 Hz at 3.8 mA. (A-C, top) Stimulation pulses (red) with overlaid ECAPs (black). (A-C, bottom) Raster plots of axonal firing and binned counts of active axons using 2-ms bins. In the raster plots, axons are ordered by the distance from the electrode, with the nearest axons shown at the bottom of the plots. At 50 Hz (A), axonal firing remained synchronized with each pulse, resulting in consistent ECAPs across the duration of stimulation. As the frequency increased to 500 Hz (B) and 1000 Hz (C), there was a reduction in firing synchrony and ECAP amplitudes decreased. (D) ECAP amplitudes for each applied stimulation pulse across tested stimulation frequencies. We calculated the ECAP amplitude following each stimulation pulse, within the time window preceding the next pulse onset. Freq. = frequency. Amp. = amplitude.

To further characterize frequency-dependent responses, we analyzed ECAP growth curves for 50, 150, 500, and 1000 Hz frequencies on a per-pulse basis (**Figure 9A-D**). For each condition, we generated growth curves for individual stimulus pulses and then averaged the ECAP amplitudes across pulses. At 50 and 150 Hz, growth curves stabilized quickly and ECAP amplitudes remained consistent across pulses (**Figure 9A-B**). In contrast, 500 and 1000 Hz stimulation produced an initial onset ECAP similar to lower frequency responses, but subsequent pulses elicited reduced and more variable ECAP amplitudes (**Figure 9C-D**). Thereafter, axons fired intermittently for subsets of pulses.

**Figure 9.**
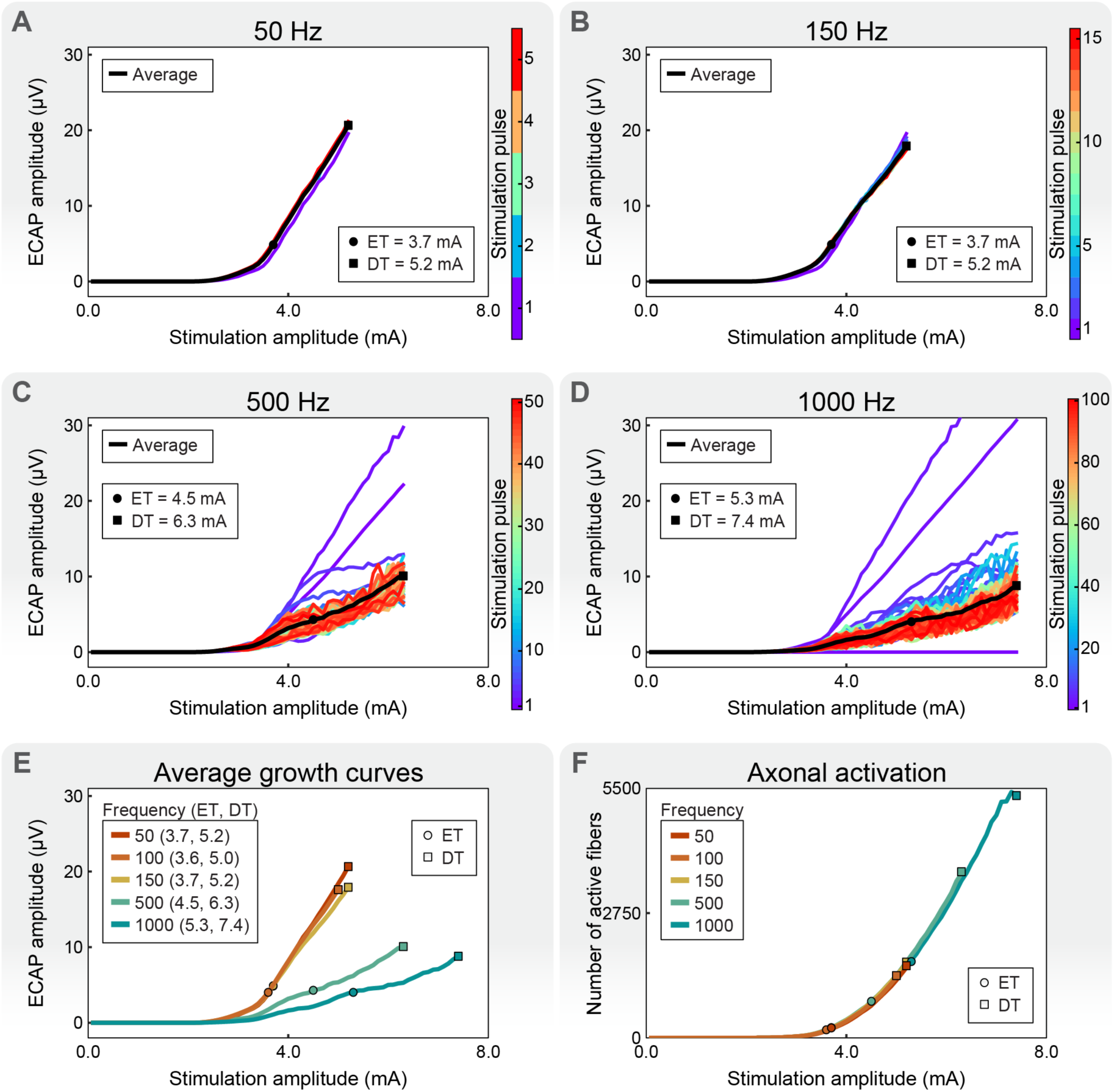
Frequency-dependent changes in evoked compound action potential (ECAP) growth curves and axonal activation. We generated growth curves for each stimulus pulse for stimulation frequencies of (A) 50 Hz, (B) 150 Hz, (C) 500 Hz, and (D) 1000 Hz. For each frequency, we assessed steady-state responses by computing the average ECAP growth curves across pulses during 100 ms of stimulation. (E) Averaged ECAP growth curves by stimulation frequency. ECAP thresholds (ETs) and discomfort thresholds (DTs) are shown in mA. (F) Counts of activated axons as a function of stimulation amplitude and frequency.

Next, we compared averaged growth curves across all frequencies (**Figure 9E**). When increasing the stimulation frequency from 50 to 150 Hz, ET remained at 3.7 mA using the averaged growth curves. However, increasing the frequency to 500 and 1000 Hz increased ETs to 4.5 mA and 5.3 mA, respectively. Finally, we evaluated the number of active fibers per stimulation amplitude across all frequencies (**Figure 9F**). Overall, the number of active fibers remained similar across conditions at equivalent stimulation amplitudes (**Figure 9F**). However, due to the higher ET and corresponding DT resulting from lower ECAP amplitudes during 1000-Hz stimulation, the number of active fibers increased by ∼246% at DT (1547 to 5353 active fibers) from 50 Hz to 1000 Hz, despite a decrease in the ECAP amplitude by ∼50% (20.7 and 8.8 µV at 50 and 1000 Hz, respectively.

## 4. Discussion

In this study we built upon prior computational modeling efforts of ECAP-based SCS to offer a systematic characterization of how relevant anatomical and technical factors affect both the underlying neural responses to SCS and corresponding ECAP recordings [11,29]. We employed a hybrid modeling approach to simulate the neural responses to SCS across a range of parameters and calculated the corresponding model ECAP recordings. To analyze each parameter, we calculated growth curves, model-predicted ETs and DTs, and neural recruitment profiles.

### 4.1. Recording configurations

Clinically, ECAPs are often recorded using differential configurations to minimize common-mode artifact [23,24,49–51,76]. However, this approach inherently reduces ECAP amplitudes [13]. Given that ECAPs are composed of propagating neural signals, their spatiotemporal properties (e.g., conduction velocity and signal dispersion) may be leveraged to optimize both ECAP amplitudes and signal-to-noise ratios in clinical applications.

We used our computational model to analyze several differential recording pairs (E1-E0, E2-E0, and E3-E0) (**Figure 2**). Using monopolar recordings, we found that the N1 peak amplitude in the differential recordings was maximized when the leading positive peak of the ECAP arrived at the distal recording contact (i.e., E0) while the negative peak arrived simultaneously at the proximal recording contact (e.g., E3). Similarly, the P2 peak amplitude was greatest when the trailing positive peak reached the proximal recording contact concurrently with the negative peak at the distal recording contact. These conditions optimized the constructive interference in the differential signal and thus resulted in the largest ECAP features.

Importantly, these effects are influenced by the spatiotemporal dispersion of the ECAP as it propagates. As the distance from the stimulation site increased, the ECAP waveform broadened, and peak amplitudes decreased due to dispersion. Because the ECAP is comprised of action potentials from axons of differing diameters, locations, and activation times, as propagation distance increases, differences in conduction velocities among axons may change both the timing and amplitude of the ECAP. This behavior also explains the observed similarities in ECAPs at ET for E2-E0 and E3-E0. At ET, fewer axons are recruited, and those that are active tend to be larger-diameters and thus exhibit faster conduction velocities. As the timing of peaks in the ECAP depend on the conduction velocity, this resulted in a shorter optimum distance between the proximal and distal recording contacts. As the stimulation amplitude increased and more axons were recruited, the average conduction velocity of the ECAP decreased and E3-E0 produced larger ECAP amplitudes than E2-E0. However, it is important to recognize that our computational model did not include sources of stimulation artifact, and utilizing electrodes in closer proximity to the stimulation site comes at the cost of the stimulation artifact encroaching on the ECAP [13,17,18,24,57]. Additionally, increasing the separation between recording contacts allows more common mode noise into the recorded signal. Consequently, the optimal distance between recording electrodes requires balancing the ability to record maximal ECAP amplitudes while minimizing artifact contamination. Overall, these findings are clinically relevant for the design and optimization of electrode arrays, particularly with respect to their spatial layout and recording configurations. In studies involving multiple electrode arrays or when recording at a sufficient distance, skipping intermediate electrodes between differential recording pairs may enhance ECAP amplitudes.

### 4.2. Anatomical considerations

Consistent with prior studies [11,15], we observed that larger dCSF thicknesses decreased ECAP amplitudes for a given stimulation amplitude, resulting in increased ETs and DTs (**Figure 3B-C**). The reduction in ECAP amplitude stemmed from the increased distance between the stimulating electrodes and spinal cord for thicker dCSF layers, which necessitated higher stimulation amplitudes to achieve equivalent axonal activation (**Figure 3D-E**). Moreover, the increased distance between the activated axons and recording electrodes reduced ECAP amplitudes, even for the same number of active axons (**Figure 3E**). Additionally, the slope of growth curves was highly sensitive to dCSF thickness (**Figure 3C**). In models with less dCSF, we found that small changes in stimulation amplitudes led to larger shifts in neural activation, and consequently ECAP amplitudes, compared to models with more dCSF. This observation suggests that clinically, the growth curve slope may be used to inform the sensitivity needed in closed-loop SCS systems, with a steeper slope indicating the need for smaller, more precise adjustments to maintain more consistent neural dosing.

At both ET and DT, the 1-mm dCSF model recruited a greater proportion of smaller-diameter axons (i.e., < 9.0 µm) than the 5-mm dCSF model (**Figure 3F**). Moreover, the cross-sectional area of activation varied considerably between the 1-mm and 5-mm dCSF models. The 5-mm dCSF model produced a larger area of activation than the 1-mm dCSF model to generate an ECAP above the 4-µV threshold for ET. Overall, these findings may provide a mechanistic explanation for altered sensory perceptions as patients change postures [68]. Furthermore, a constant ECAP amplitude does not necessarily reflect equivalent underlying neural activation [11].

### 4.3. Stimulation parameters: pulse width

Prior studies have demonstrated that pulse width significantly influences ECAP characteristics, with longer pulse widths increasing ECAP latency [11,12]. Our findings are consistent with these observations and further elucidate how pulse width influences neural recruitment dynamics and ECAP characteristics. As expected, increasing pulse width reduced ET and DT and increased ECAP latency (**Figure 4B-C**).

Despite similar ECAP amplitudes at ET across pulse widths, there were differences in the underlying neural recruitment. Consistent with previous computational modeling, we observed that longer pulse widths preferentially recruited a larger proportion of smaller-diameter fibers (**Figure 4F**) [30]. Specifically, at ET, stimulation with both 90-µs and 300-µs generated ECAPs that were primarily composed of activation of axons with diameters in the range of 10.0-11.9 µm (**Figure 4E**). However, stimulation with the 300-µs pulse width recruited additional axons in the diameter range of 8.0-9.9 µm, which produced an additional peak following P2 (**Figure 4E, bottom row**) [12]. These results highlight that comparable ECAP amplitudes may mask differences in the number, timing and underlying population of axons recruited. Additionally, when optimizing pulse width in closed-loop, ECAP-based SCS systems, it is important to account for how changes in pulse width can affect recruitment dynamics, and ECAP timing and amplitude.

### 4.4. Stimulation parameters: interphase interval

We found that increasing the IPI from 0 to 90 µs decreased activation thresholds and lowered both ET and DT, with no further changes for IPIs longer than 90 µs (**Figure 5B-E**). At ET, the 120-µs IPI activated axons during both phases of the symmetric biphasic stimulation pulse (**Figure 5F, bottom left**). The first phase of stimulation primarily recruited axons near E7, while the second phase activated axons near E6. This temporal and spatial offset caused interference in the total recorded ECAP (**Figure 5F, bottom right**). In contrast, when stimulating with a 0-µs IPI at ET, the second phase recruited relatively fewer axons (**Figure 5F, top**). As a result, although the 120-µs IPI activated axons more efficiently, it required the largest number of axons to generate a given ECAP amplitude.

Our findings highlight a complex interaction between stimulation waveform parameters and electrode configurations in shaping recorded ECAPs. The spatial and temporal offset of activation introduced by longer IPIs, particularly when utilizing bipolar stimulation configurations, can cause interference in the total recorded ECAP. Consistent with our observations with increased pulse widths, we found that longer IPIs introduced a latency shift in ECAPs.

### 4.5. Stimulation parameters: waveform

Among tested waveforms, rectangular-biphasic stimulation was the most efficient (**Figure 6B-C**). Triphasic stimulation had higher ET and DT values and produced delayed ECAPs as neural activation was primarily driven by the second phase of stimulation. Although this latency shift may aid in the mitigation of stimulation artifact, it comes at the expense of reduced stimulation efficiency relative to rectangular-biphasic stimulation.

Overall, our results underscore the importance in the selection of waveform morphology and polarity on neural recruitment and ECAP features during SCS. While asymmetric waveforms may offer benefits, such as artifact reduction or altered neural activation, they may also compromise stimulation efficiency and reduce ECAP fidelity [13,14,17]. Additionally, our findings are consistent with previous work suggesting that phase asymmetry can introduce complex activation patterns and ECAP morphologies [17,79].

### 4.6. Stimulation parameters: stimulation configuration

Guarded-cathode stimulation was most efficient with the lowest thresholds and largest ECAP amplitudes across configurations (**Figure 7B-C**). Additionally, guarded-cathode stimulation was associated with shorter ECAP latencies than bipolar, wide-bipolar, and guarded-anode stimulation (**Figure 7B**). This effect was due to the activation which occurred during the first phase of stimulation near E6. While ECAPs from bipolar and wide bipolar stimulation were similarly composed from activation during the first phase of stimulation, activation for these stimulation configurations occurred near E7. As E7 was further from the recording site than E6, this increased the propagation distance and delayed the recorded ECAPs.

Our findings highlight that the choice of stimulation configuration must account for both the stimulation electrode positioning relative to the recording electrodes and the timing of neural activation. When using two leads or when stimulation and recording electrodes are separated by a sufficient distance, our computational modeling suggests that guarded-cathode stimulation is most efficient because of the more focused electric field compared to bipolar stimulation. However, when stimulation and recording occur on the same lead or across shorter distances, it is critical to increase spatial separation to allow the neural signal to propagate beyond the stimulation artifact. In such cases, bipolar configurations with the cathode located at the most distal contact to the recording pair may improve recording quality.

### 4.7 Stimulation parameters: stimulation frequency

To better understand how stimulation frequency influences ECAP recordings and neural responses during SCS, we used our computational modeling framework to examine the impact across frequencies from 50 to 1000 Hz. Increasing stimulation frequency altered both ECAP characteristics and the underlying axonal firing dynamics (**Figure 8**). At conventional SCS frequencies (i.e., 50 to 150 Hz), axons exhibited one-to-one entrainment with applied stimulation pulses, resulting in stable ECAP amplitudes and growth curves (**Figure 8A, D**; **Figure 9A-B**). In contrast, during high-frequency stimulation (i.e., 500 and 1000 Hz), the intervals between stimulation pulses are shorter than the axon’s refractory periods. As a result, some axons fail to fire in a one-to-one manner with each stimulation pulse (**Figure 8B-D**). This partial entrainment reduced firing synchrony across the population of axons and is consistent with overdrive desynchronization [21].

The number of active fibers remained similar across frequencies at equivalent stimulation amplitudes (**Figure 9F**). However, ETs increased when stimulation was applied at 500 and 1000 Hz when growth curves were averaged across stimulation pulses (**Figure 9E**). Together, these results indicate that elevated ETs at increased stimulation frequencies reflect reduced firing synchrony across stimulation pulses rather than changes in the activated fiber populations. As axons fail to fire for every stimulation pulse, action potentials become temporally dispersed, thus reducing the synchronous summation that produces the ECAP and increasing the stimulation amplitude required to reach ET. Overall, our findings suggest that ECAP amplitudes during high-frequency SCS are strongly influenced by frequency-dependent neural firing dynamics.

### 4.8 Limitations

While the computational framework presented in our study offers valuable insights into neural activation and ECAP characteristics during SCS, several limitations should be considered when interpreting the results. First, our model ECAP recordings do not incorporate sources of electrical or biological noise. In practice, ECAPs may be contaminated by stimulation artifact, movement artifact, respiratory signals, or electromyographic (EMG) activity [12,13,17,23,25]. These sources of artifact can obscure true neural signals and complicate the interpretation of ECAPs. Notably, EMG activity may propagate across recording electrodes and be misinterpreted as a neural signal [17]. While this simplification reduces the physiological realism of our models, it enables the isolation and analysis of the direct neural response to stimulation without interference from non-neural signal components. Second, our models did not include dorsal rootlets and future modeling efforts should consider these fibers as well as how variations in axon trajectories and morphologies influence neural responses and ECAPs. Finally, our FEM model was constructed using averaged anatomical and conductivity values from the literature. While canonical models have proven useful in exploring the mechanisms of SCS [15,29,31,80,81], they do not capture variability in anatomy, tissue properties, or electrode placement [82,83]. Future work should incorporate patient-specific FEM models derived from imaging data and directly compare model predictions with clinical recordings to enhance model validity and provide further mechanistic insights.

## 5 Conclusions

In this study, we used a computational modeling framework to systematically examine how anatomical variability, stimulation parameters, and recording configurations influence neural recruitment and ECAP characteristics during SCS. We found that factors such as dCSF thickness significantly affect activation thresholds, recruitment patterns, and ECAP morphology. Additionally, our results demonstrate that by leveraging the propagation dynamics of ECAPs, the selection of recording electrodes can be chosen to maximize ECAP amplitudes. Stimulation parameters introduced trade-offs between stimulation efficiency, axon selectivity, and ECAP timing and morphology. Notably, similar ECAP amplitudes may mask differences in underlying neural activation, highlighting the importance of interpreting ECAPs within their anatomical and technical context. Our findings provide insights into how ECAPs may encode specific aspects of neural activation, supporting their utility as real-time feedback signals and providing a mechanistic foundation for optimizing closed-loop SCS systems.

## Acknowledgments

This research was supported by a research grant from Medtronic, Inc., the Rackham Merit Fellowship Program at the University of Michigan, and computational resources and services provided by Advanced Research Computing at the University of Michigan.

---

**Supplementary Figure 1:**
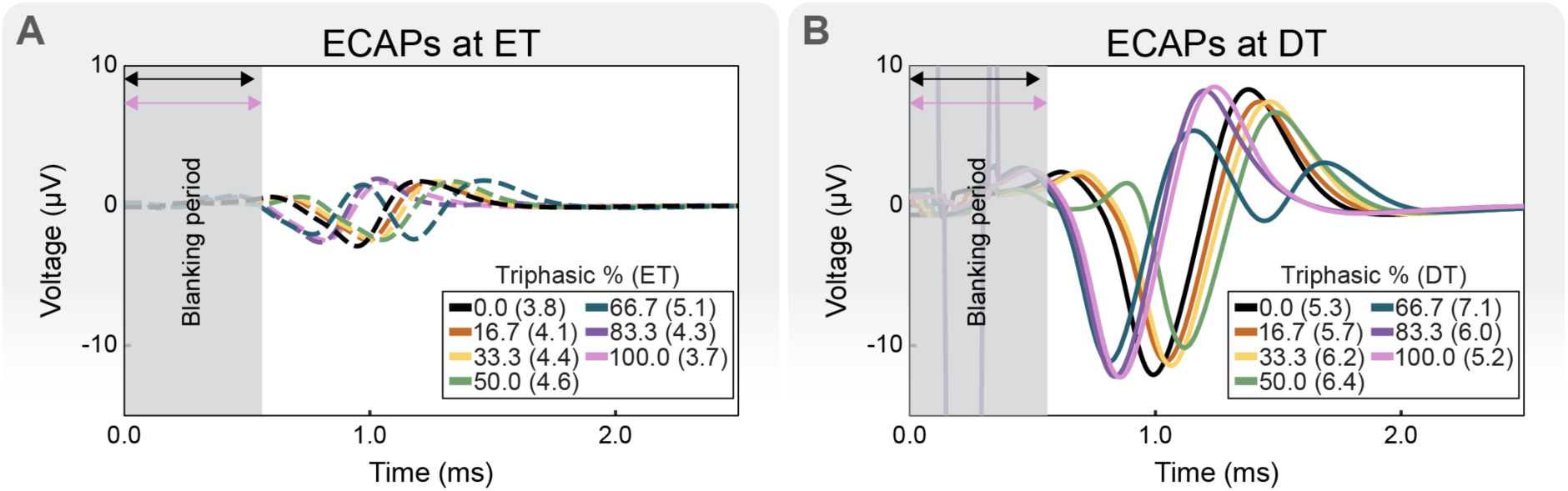
Evoked compound action potential (ECAP) recordings using triphasic waveforms and asymmetric phase durations. (A) ECAPs at ECAP threshold (ET). (B) ECAPs at discomfort threshold (DT). ETs and DTs are reported in mA.

